# STAT3 and GR cooperate to drive basal-like triple negative breast cancer gene expression and proliferation

**DOI:** 10.1101/730606

**Authors:** Megan E. Conway, Joy M. McDaniel, James M. Graham, Katrin P. Guillen, Patsy G. Oliver, Stephanie L. Parker, Peibin Yue, James Turkson, Donald J. Buchsbaum, Bryan E. Welm, Richard M. Myers, Katherine E. Varley

## Abstract

Breast cancers can be divided into subtypes with different prognoses and treatment responses based on global gene expression differences. Luminal breast cancer gene expression and proliferation are driven by the transcription factors Estrogen Receptor α (ER), FOXA1 and GATA3. Targeting ER is the most effective therapy for treating luminal breast cancer because ER is the master regulator of the luminal gene expression program. In contrast, it is unclear which transcription factors are responsible for driving the gene expression signature that defines basal-like triple negative breast cancer, and there are no targeted therapies approved to treat this aggressive subtype of the disease. This study utilized integrated analysis of DNA methylation, chromatin accessibility, transcription factor binding, and gene expression in large collections of breast cancer cell lines and patient tumors to identify transcription factors responsible for the basal-like gene expression program. The results of this study indicate that glucocorticoid receptor (GR) and signal transducer and activator of transcription 3 (STAT3) bind to the same genomic regulatory regions that are specifically open and unmethylated in basal-like breast cancer. These transcription factors cooperate to regulate expression of hundreds of genes in the basal-like gene expression signature and these downstream genes are associated with poor prognosis in patients. Furthermore, combination treatment with small molecule drugs that inhibit both transcription factors leads to synergistic decreases in cell proliferation in cell lines and patient-derived organoid models. This study demonstrates that GR and STAT3 cooperate to regulate the basal-like breast cancer gene expression program and provides the basis for improved therapy for basal-like triple negative breast cancer through rational combination of STAT3 and GR inhibitors.

## Introduction

Breast cancers can be divided into subtypes with different prognoses and treatment responses based on global gene expression differences(1,2). These expression differences reflect the transcription factors (TFs), epigenetic states and gene regulatory networks in the normal cells from which the tumors arise, as well as changes that accumulate during tumorigenesis.

Normal luminal breast cells proliferate in response to estrogen through activation of the TF estrogen receptor α (ER) and ER’s downstream target genes. Breast cancers that express ER are most often classified into the luminal subtype and likely arise from luminal breast cells(1). These luminal ER+ tumors are treated with hormone therapies that prevent ER from activating its pro-proliferative transcriptional program. Even though these tumors usually have wildtype ER, this is an extremely effective targeted therapy because it inhibits the master regulator of gene expression and proliferation in this cell type.

In contrast, 15-20% of breast cancer patients are diagnosed with triple negative breast cancer (TNBC), which is defined by a lack of expression of ER and progesterone receptor, and normal expression of human epidermal growth factor receptor 2. There are no FDA approved targeted therapies to treat TNBC. Even after aggressive treatment with surgery, radiation and cytotoxic chemotherapy, TNBC patients have a high rate of recurrence within 3 years (3), and a high rate of death from disease (38%) (4). There is a crucial need for alternative therapeutic strategies for TNBC. Approximately 70-80% of TNBC tumors are classified into the basal-like breast cancer gene expression subtype, and likely arise from myoepithelial breast cells(5). It is unclear which TFs are responsible for regulating the pro-proliferative transcriptional program in the basal-like TNBC subtype. The goal of this study is to identify TFs that regulate clinically relevant gene regulatory programs in basal-like breast cancer, and determine if inhibiting these TFs could provide novel therapeutic strategies to improve clinical outcomes for basal-like TNBC patients.

The role of many TFs have been studied in basal-like breast cancer, including oncogenic drivers such as MYC (6), and other TFs that regulate pathways associated with basal-like phenotypes such as EN1 (7), GLI1 (8), NF-kB (9), HIF1α (10), TWIST1 (11), FOXC1 (12,13), AP-1 (14), GR(15-18), STAT3(19,20), and MAFK (21). Many of these TFs regulate important pathways in basal-like breast cancer and these TFs often regulate each other’s expression (22,23). Several of these TFs can be inhibited with small molecule drugs and represent promising therapeutic targets(18,24,25). What remains unclear is which TFs are responsible for driving the gene expression signature that defines the basal-like TNBC subtype and which are downstream of the master regulator.

This study utilized integrated analysis of DNA methylation, chromatin accessibility, TF binding, gene expression, and proliferation in large collections of breast cancer cell lines and patient tumors to identify TFs that drive the basal-like TNBC gene expression program. We sought to identify regulatory regions that are specifically unmethylated and have accessible chromatin in basal-like TNBC compared luminal breast cancers. We identified TFs whose motifs and binding sites are most highly enriched at these basal-specific regulatory regions. We then investigated whether these TFs were more highly expressed in basal-like TNBC compared to luminal breast cancer, whether the TFs regulated genes in the basal-like TNBC gene expression program, and whether the genes regulated by these TFs were associated with patient outcomes. This genome-wide approach led to the discovery that cooperation between two TFs (STAT3 and GR), rather than a single ‘master regulator’, drives expression of hundreds of genes in the basal-like TNBC gene expression program. This TF cooperation provides basal-like TNBC cells with their resilient proliferative capacity. Furthermore, inhibiting both TFs simultaneously led to synergistic decreases in cell proliferation in basal-like TNBC cell lines and patient-derived organoid models, which suggests a new treatment strategy for basal-like TNBC.

## Results

### Identification of regulatory regions and candidate TFs specific to basal-like breast cancer

We hypothesized that genomic regulatory regions that are specifically unmethylated in basal-like breast cancer compared to luminal breast cancer will be enriched for TF binding sites involved in controlling basal-specific gene expression. To identify regulatory regions specific to basal-like breast cancer, reduced representation bisulfite sequencing (RRBS) was performed on 28 breast cancer cell lines (18 basal-like and 10 luminal) in order to measure DNA methylation across the genome. Of the 479,746 CG positions in the genome with at least 10x coverage in each cell line, 3,748 CGs were significantly differentially methylated (Linear Regression Benjamini-Hochberg adjusted p< 0.05) between basal-like and luminal cell lines **(Figure 1A)**.

**Figure 1.**
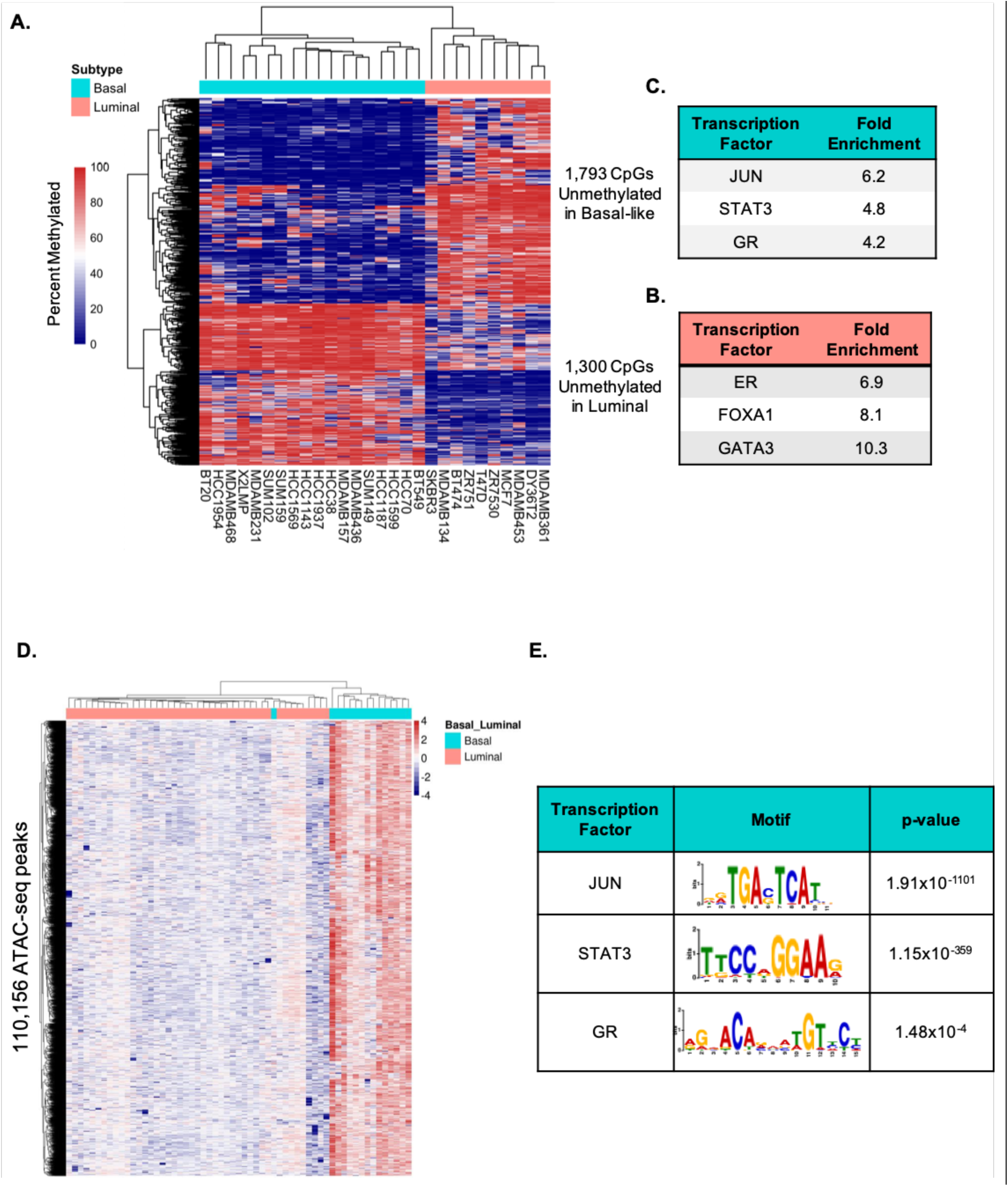
Identification of candidate TFs enriched at regions that are unmethylated and have open chromatin in basal-like compared to luminal breast cancer**. A.** Heatmap of CG positions that are significantly differentially methylated between basal-like and luminal breast cancer cell lines (Linear Regression Benjamini-Hochberg adjusted p< 0.05). Enrichment of TF binding sites from ChIP-seq experiments in a variety of cell lines that overlap CG positions that are specifically unmethylated in luminal (**B**) and basal-like (**C**) cell lines. **D.** Heatmap of ATAC-seq peaks that have significantly more open chromatin in basal-like patient tumors compared to luminal tumors (DESEQ2 Adjusted p<0.05). **E.** Enrichment for JUN, STAT3, and GR motifs in the basal-specific ATAC-seq peaks.

There were 1,300 CGs in intergenic or promoter regions of the genome that were significantly unmethylated in luminal breast cancer. These CG positions were intersected with the locations of TF binding sites from ChIP-seq experiments performed on 149 TFs by the ENCODE Project in a variety of cell lines (26). To determine which TFs were enriched at regions that are specifically unmethylated in luminal breast cancer cells, the fraction of binding sites for each TF that intersect these CGs was divided by the fraction of binding sites for each TF in the ENCODE dataset. The most highly enriched sequence-specific TF binding sites were ER (6.9 fold), FOXA1 (8.1 fold), and GATA3 (10.3 fold). This confirms that this approach is sound because ER, FOXA1, and GATA3 are known to be the master regulators of the luminal gene expression program (27,28) **(Figure 1B)**.

There were 1,793 CGs that were in intergenic or promoter regions of the genome and were specifically unmethylated in basal-like breast cancer. These CG positions were intersected with the TF binding sites locations from ChIP-seq experiments performed on 149 TFs by the ENCODE Project (26). The most highly enriched sequence-specific TFs in these regions were JUN (6.2 fold), STAT3 (4.8 fold), and GR (4.2 fold) **(Figure 1C)**. This result led us to hypothesize that JUN, STAT3, and GR are involved in regulating basal-like breast cancer gene expression.

To further investigate this hypothesis, we sought to use primary human tumors rather than cell lines, to directly investigate chromatin accessibility with ATAC-seq rather than inferring it from a lack of DNA methylation, and use TF motif enrichment rather than the ENCODE Project ChIP-seq experiments, which did not include any basal-like breast cancer cell lines. We analyzed ATAC-seq data from 59 primary breast tumors (15 basal-like and 44 luminal)(29). There were 110,156 regions of the genome that exhibit open chromatin specifically in basal-like tumors compared to luminal tumors (DESEQ2 Adjusted p<0.05) (**Figure 1D**). Motif analysis found significant enrichment for the canonical motifs for JUN/AP1 (p=1.91×10^-1101^), STAT3 (p=1.15×10^-359^), and GR (p=1.15×10^-4^) in the chromatin regions that were specifically open in basal-like tumors and closed in luminal tumors (**Figure 1E**). While the enrichment for the GR motif is significant, it is lower than that of STAT3 and JUN/AP1, which is consistent with previous studies that suggest GR can be tethered to some enhancers through protein:protein interactions, rather than direct DNA binding(30).

### Expression of GR and STAT3 is higher in basal-like breast cancer and associated with a worse prognosis

If JUN, GR and STAT3 regulate the basal-like breast cancer gene expression program, then one would expect these TFs to have higher expression in basal-like breast cancer compared to luminal breast cancer. RNA-seq was performed on the same 28 breast cancer cell lines used for DNA methylation analysis. Both GR and STAT3 have significantly higher expression in basal-like cell lines compared to luminal cell lines, while JUN does not (**Figure 2A**). An independent gene expression dataset containing 19 basal-like breast cancer cell lines and 21 luminal breast cancer cell lines (31) confirmed that both GR and STAT3 have higher expression in basal-like breast cancer (**Figure 2B**). These results confirm that GR and STAT3 have higher expression in basal-like breast cancer than luminal breast cancer.

**Figure 2.**
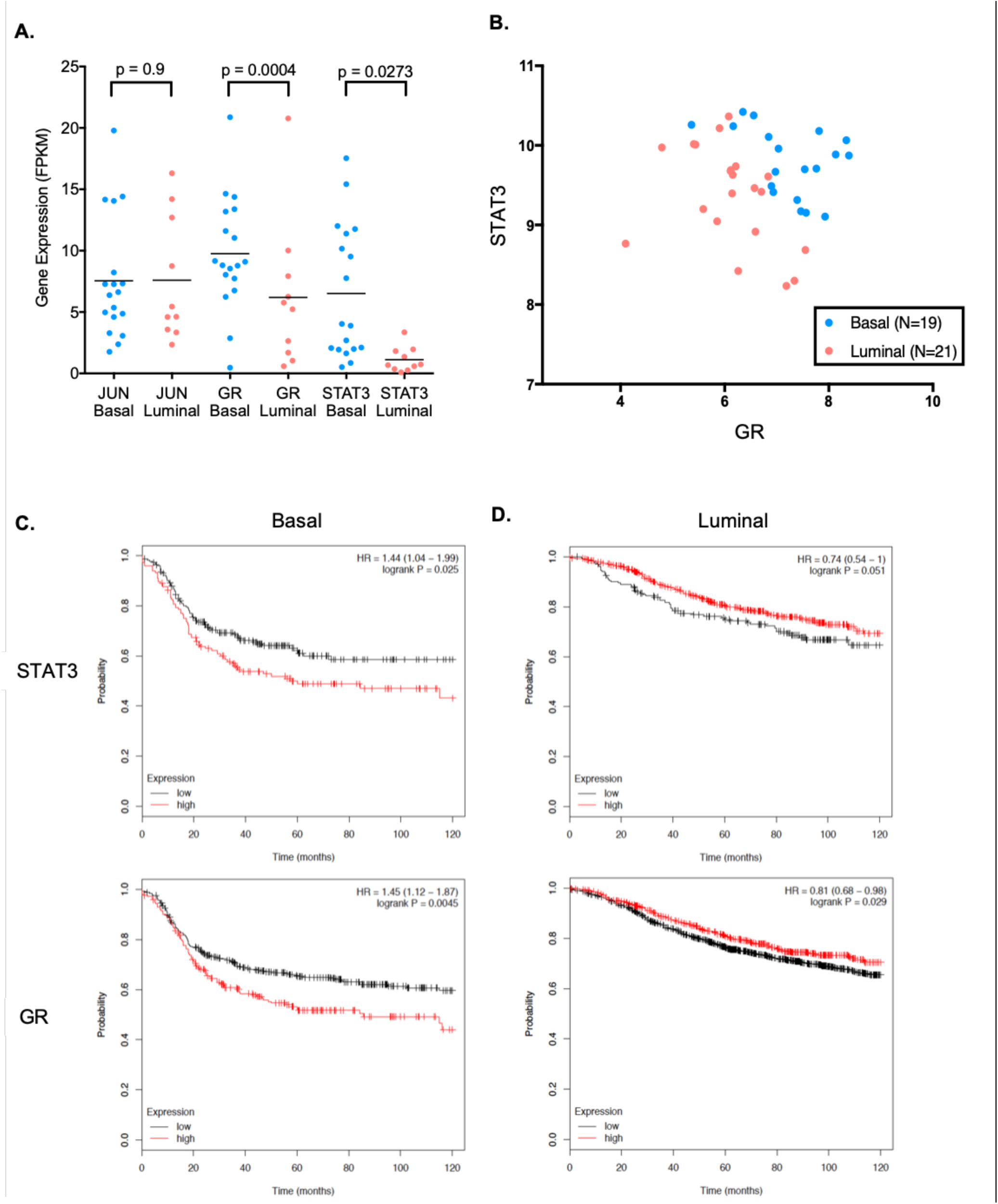
Expression of GR and STAT3 is higher in basal-like breast cancer and associated with shorter relapse-free survival. **A.** RNA-seq data shows GR and STAT3 have higher expression in basal-like (N=18) compared to luminal (N=10) cell lines, while JUN does not. Mann Whitney p-values displayed. **B.** An independent microarray gene expression dataset shows GR and STAT3 have higher expression in basal-like (N=19) compared to luminal (N=21) cell lines. **C.** In basal-like breast cancer patients, higher expression of GR and STAT3 is associated with shorter relapse-free survival. **D**. In contrast, luminal breast cancer patients have better prognosis when GR and STAT3 are highly expressed.

Some luminal breast cancers do express low levels of GR and STAT3, so we investigated the association between prognosis and GR or STAT3 expression. Analysis of a dataset containing breast tumor gene expression and long-term survival data (32) revealed that higher expression of both GR and STAT3 was associated with shorter relapse-free survival in basal-like breast cancer (**Figure 2C**). In contrast, luminal breast cancer patients with higher GR and STAT3 expression were more likely to have longer relapse-free survival (**Figure 2D**). These results confirm that GR and STAT3 expression has a unique role in basal-like breast cancer and suggest that these TFs are associated with aggressive features of the disease.

### GR and STAT3 bind to the same basal-specific regulatory regions across the genome

To directly test the hypothesis that GR and STAT3 bind to the genome and regulate the basal-like gene expression program, ChIP-seq was performed for GR and STAT3 in 8 breast cancer cell lines (4 basal-like, 4 luminal). GR is a steroid hormone nuclear receptor that is chaperoned in the cytoplasm until it binds glucocorticoid ligands which allow it to homo-dimerize and translocate to the nucleus and regulate gene expression. To assess the binding of GR, cell lines were treated with 100nM dexamethasone for 1 hour prior to performing ChIP-seq. STAT3 is a TF whose dimerization and translocation to the nucleus is controlled through phosphorylation by Janus kinases (JAK). Normally, JAK phosphorylates STAT3 when cytokine receptors bind ligands such as interferons, epidermal growth factor, and interleukin 5 and 6. STAT3 is constitutively phosphorylated and active in the majority of basal-like cell lines and patient tumors (19,33). Thus, STAT3 ChIP-seq was performed on cells in standard media.

The genome-wide GR and STAT3 binding sites from ChIP-seq were used to perform unsupervised hierarchical clustering of the cell lines based on Spearman correlations. This analysis revealed that the basal-like and luminal subtypes have distinct genome-wide GR and STAT3 binding profiles (**Figure 3A**). The basal-like cell lines form two distinct sub-clusters within the basal-like group, which corresponds to previous classifications as Basal A (HCC70 and HCC1937) and Basal B (SUM159 and MDA-MB-231) subtypes (34). Despite these differences, the four basal-like cell lines are still more similar to each other than they are to luminal cell lines when considering genome-wide GR and STAT3 binding. DESEQ2 was used to identify regions of the genome that exhibited subtype-specific binding of GR or STAT3. There were 2,002 regions of the genome that were bound by GR specifically in basal-like breast cancer (DESEQ2 adjusted p<0.05) (**Figure 3B**), and 3,667 regions of the genome that were bound by STAT3 specifically in basal-like breast cancer (DESEQ2 adjusted p<0.05) (**Figure 3C**). This result demonstrates that both GR and STAT3 bind thousands of regions of the genome in basal-like breast cancer that they do not bind in luminal breast cancer, which supports the hypothesis that these TFs regulate the basal-like gene expression program.

**Figure 3.**
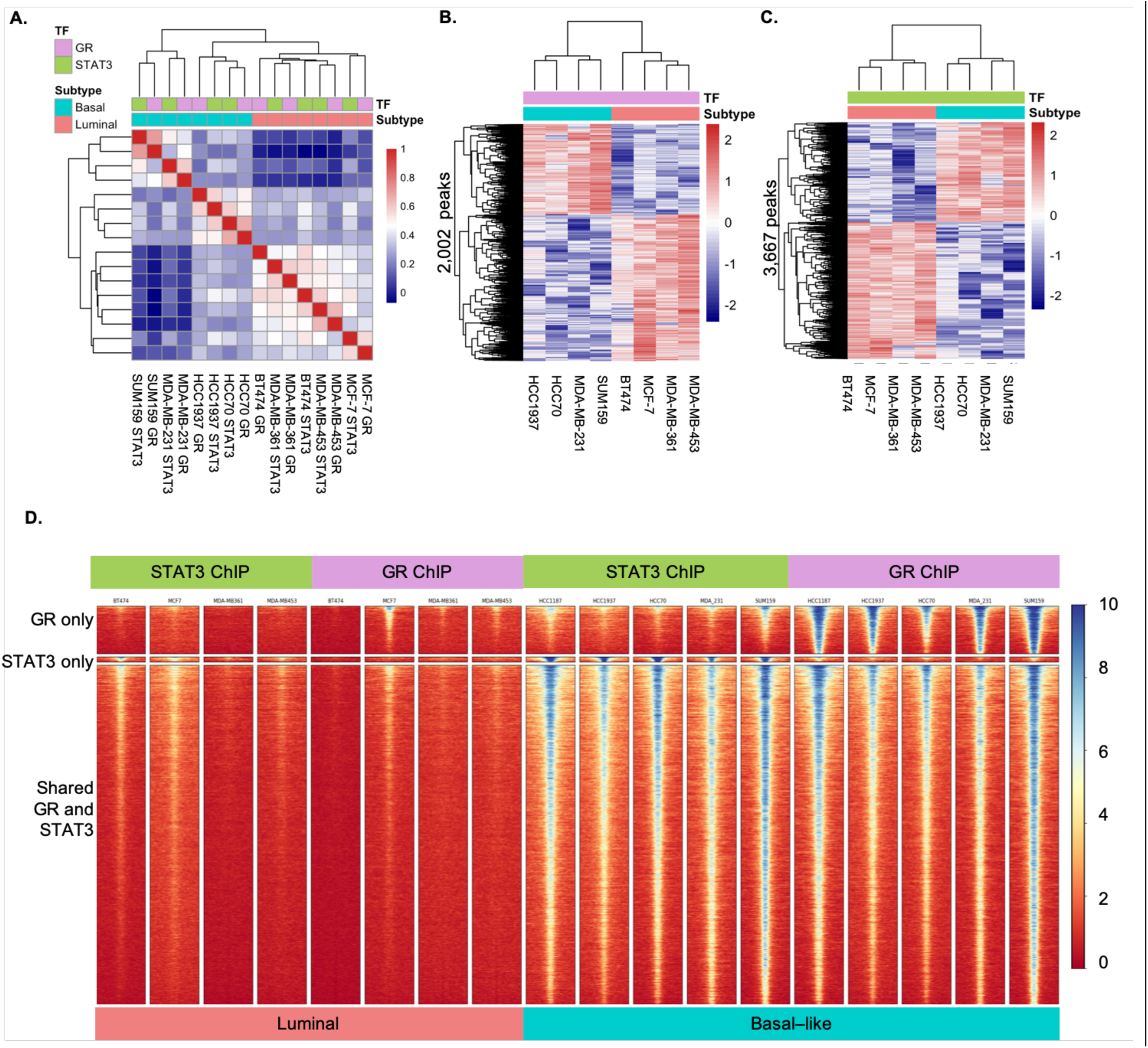
ChIP-seq of GR and STAT3 in basal-like (N=4) and luminal (N=4) cell lines. **A.** Pairwise correlations of genome-wide binding across basal-like and luminal cell lines show that GR and STAT3 exhibit global subtype-specific binding. **B.** GR exhibited subtype-specific binding at 2,002 peaks (DESEQ2 adjusted p<0.05). **C.** STAT3 exhibited subtype-specific binding at 3,667 peaks (DESEQ2 adjusted p<0.05). **D.** There are 12,712 sites bound by both GR and STAT3 specifically in basal-like cell lines.

The correlation heatmap revealed that genome-wide GR binding sites and STAT3 binding sites were highly correlated within each of the four basal-like cell lines. To investigate this further, DESEQ2 was used to perform a multivariate analysis that combined both GR and STAT3 ChIP-seq datasets to identify sites where these TFs bound alone or together in a basal-like specific manner. This analysis revealed 1,773 sites were bound by GR alone, 188 sites were bound by STAT3 alone, and 12,712 ChIP-seq peaks with both GR and STAT3 binding specifically in basal-like cell lines (**Figure 3D**). This result indicates that GR and STAT3 bind the same basal-specific regulatory regions more often than they bind alone.

The majority of shared GR and STAT3 basal-like specific binding sites are 50-500 kb from the nearest gene transcription start site, indicating that they are likely binding distal regulatory elements such as enhancers (**Figure 4A**). Motif analysis was performed on the shared GR and STAT3 basal-like specific binding sites. The canonical STAT3 motif was found in 5,044 of these peaks, resulting in an E-value of 4.97×10^-425^ compared to shuffled control sequences. The canonical GR motif was found in 1,309 of the peaks, resulting in an E-value of 8.00×10^-37^ compared to shuffled control sequences. This result indicates that GR and STAT3 shared binding sites are significantly enriched for motifs recognized by both TFs.

**Figure 4.**
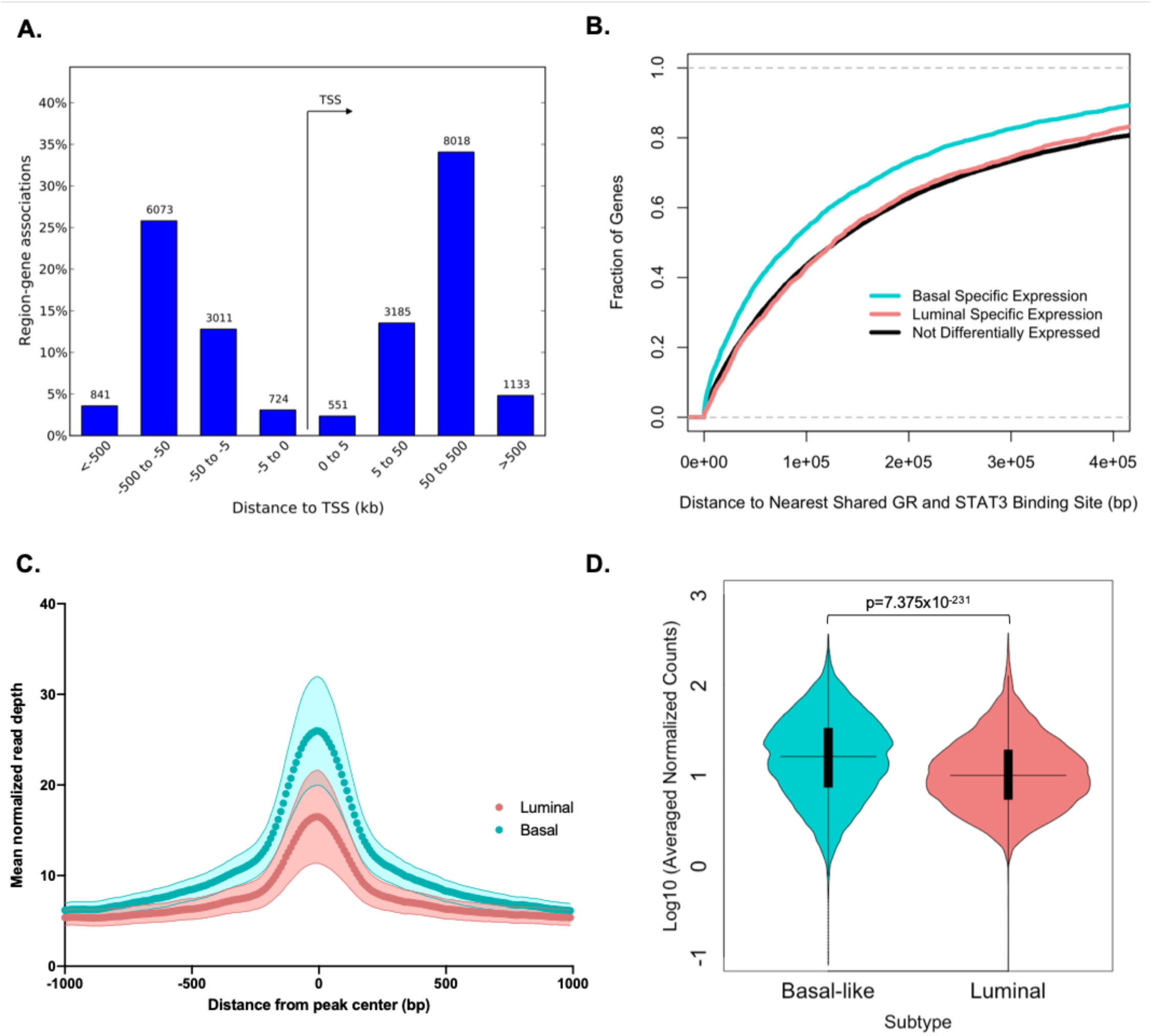
A. Basal-specific shared GR and STAT3 binding sites are most often 50-500 kb from transcription start sites (TSS) of genes. **B.** Basal-specific shared GR and STAT3 sites are closer to genes that are expressed specifically in basal-like breast cancer compared to genes expressed in luminal breast cancer and genes that are not differentially expressed between subtypes. **C.** Basal-specific shared GR and STAT3 sites have more accessible chromatin in ATAC-seq data from basal-like patient tumors (N=15) compared to luminal patient tumors (N=44) when averaged across sites. Mean (dark points) and standard deviation (lighter band) of averaged normalized read counts across patients are depicted. **D.** Violin plot of ATAC-seq data for 12,712 Basal-specific shared GR and STAT3 shows sites have more accessible chromatin in basal-like patient tumors (N=15) compared to luminal patient tumors (N=44). Mann Whitney p-value, median (horizontal line), interquartile range (black box), and range (vertical line) are depicted.

To test the hypothesis that shared GR and STAT3 basal-specific binding sites are near genes that are specifically expressed in basal-like breast cancer, RNA-seq data from 59 primary breast tumors (15 basal-like and 42 luminal) were analyzed(35). There were 7,121 genes that had significantly higher expression in basal-like breast tumors, and 3,464 genes that had significantly higher expression in luminal breast tumors (DESEQ adjusted p-value 0.05). The distance between the transcription start site of each gene and the nearest shared GR and STAT3 binding site was calculated. Genes that had significantly higher expression in basal-like breast tumors were closer to shared GR and STAT3 basal-specific binding sites than genes that were higher in luminal tumors or genes that were not differentially expressed (**Figure 4B**).

To further investigate whether the shared GR and STAT3 basal-specific binding sites discovered in cell lines are open and accessible in patient tumors, analysis of ATAC-seq data from 57 primary breast tumors (15 basal-like and 44 luminal) was performed (29). The shared GR and STAT3 basal-specific binding sites are significantly more open in basal-like patient tumors compared to luminal patient tumors (Mann Whitney p=7.375×10^-231^) (**Figure 4C & 4D**).

Together these results indicate that GR and STAT3 bind together at thousands of regulatory regions that are specifically open in basal-like tumors and closed in luminal tumors, and these binding sites are near genes that have significantly higher expression in basal-like tumors. These data further support the hypothesis that GR and STAT3 regulate the basal-like gene expression program.

### GR and STAT3 cooperatively regulate gene expression

To determine which genes are regulated by GR and STAT3, RNA-seq was performed on two basal-like cell lines (HCC70 and MDA-MB-231) treated with 100nM dexamethasone or an equal volume of 100% ethanol vehicle control for 4 hours, and STAT3 siRNA SMARTpool or Non-targeting siRNA for 96 hours. Dexamethasone induction of GR resulted in significant changes in the expression of 546 genes in both cell lines (DESEQ2 Adjusted p<0.05)(**Figure 5A**). STAT3 siRNA knockdown resulted in significant changes in the expression of 1,967 genes in both cell lines (DESEQ2 Adjusted p<0.05)(**Figure 5B**).

**Figure 5.**
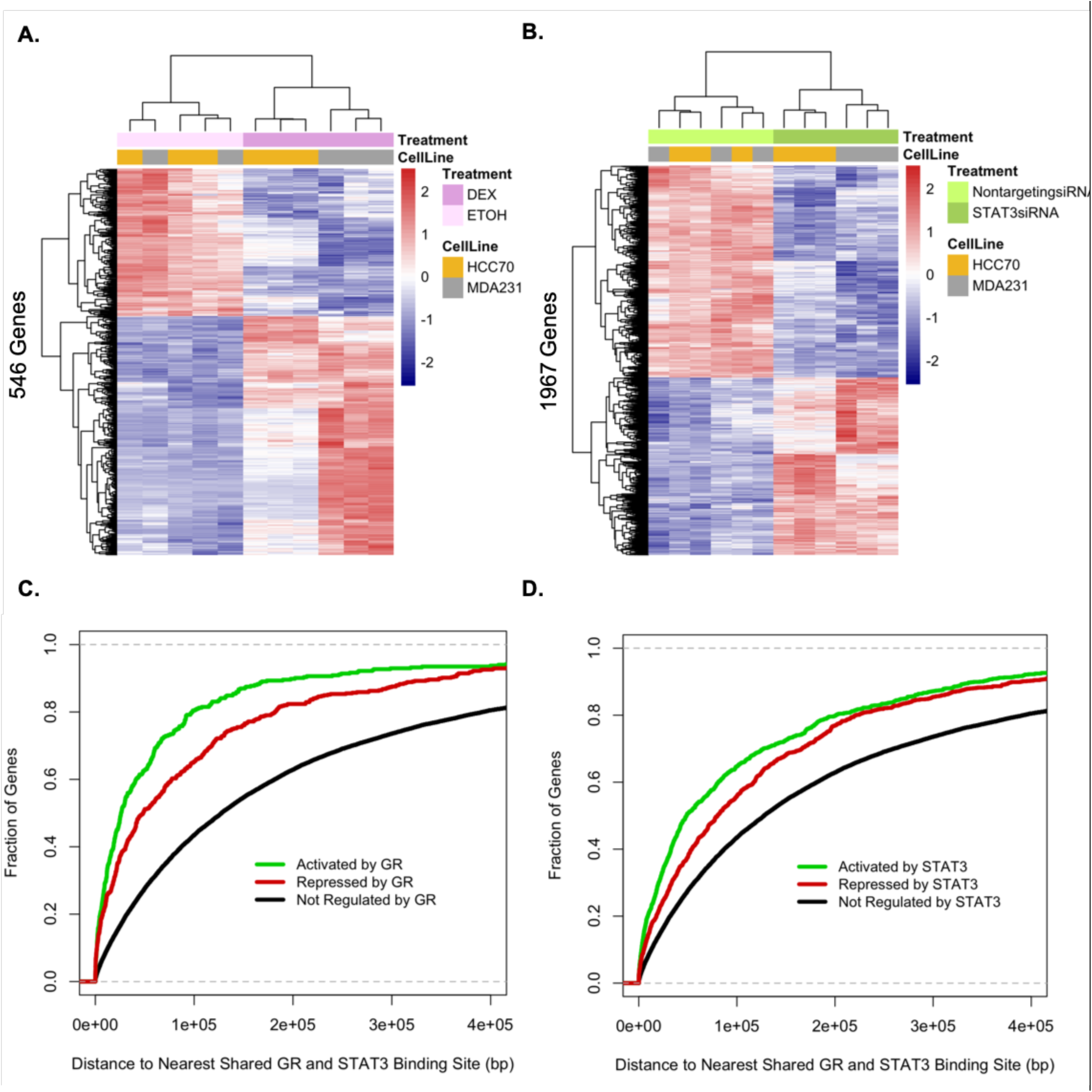
A. Treatment with dexamethasone to induce GR activity causes 546 genes to significantly change expression in two basal-like breast cancer cell lines (HCC70 and MDA-MB-231) (DESEQ2 Adjusted p<0.05). **B.** Knockdown of STAT3 with siRNA causes 1967 genes to change expression in two basal-like breast cancer cell lines (DESEQ2 Adjusted p<0.05). **C&D.** The genes that change expression when GR or STAT3 are modulated are closer to basal-specific shared GR and STAT3 binding sites than genes that do not change expression.

To test the hypothesis that genes that change expression are near the shared GR and STAT3 basal-like specific binding sites, the distance between the transcription start site of each gene and the nearest shared GR and STAT3 binding site was calculated. Previous studies have reported these TFs are more likely to activate gene expression directly, and that associated repression is more likely a downstream indirect effect(36). Therefore, the genes that are activated and repressed by each TF were analyzed separately. This analysis revealed that a larger fraction of genes that are activated by GR or STAT3 are close to a shared site when compared to repressed genes, or genes that did not change expression when these TFs were modulated (**Figure 5C & 5D**). Notably, 80% of genes that were activated by dexamethasone induction of GR and 60% of genes that are activated by STAT3 are within 100,000 bp of a shared GR and STAT3 binding site.

To further elucidate how GR and STAT3 cooperate to regulate gene expression, the MDA-MB-231 cell line was treated with dexamethasone (1µM for 24 hours) and the STAT3 inhibitor SH4-54 (8µM for 24 hours) (24) alone and in combination. A multivariate linear model with a GR:STAT3 interaction term was used to identify genes that are differentially expressed when both TFs are fully active in the nucleus. There were 970 genes that had significantly higher expression when both GR and STAT3 were active in the nucleus (BH adjusted p-value<0.05) (**Figure 6A**). Strikingly, the majority of these genes (769) fit a synergistic model of activation, where the presence of both GR and STAT3 in the nucleus resulted in higher expression than the sum of the expression from GR and STAT3 alone. Additionally, 201 genes fit an additive model, where GR and STAT3 each contribute additively to the expression. There were also 628 genes that had significantly lower expression when both GR and STAT3 were active in the nucleus (BH adjusted p-value<0.05) (**Figure 6A**). Again, the majority (515 genes) fit a synergistic model, where the presence of both GR and STAT3 in the nucleus resulted in greater repression than the sum of the repression from GR and STAT3 alone.

**Figure 6.**
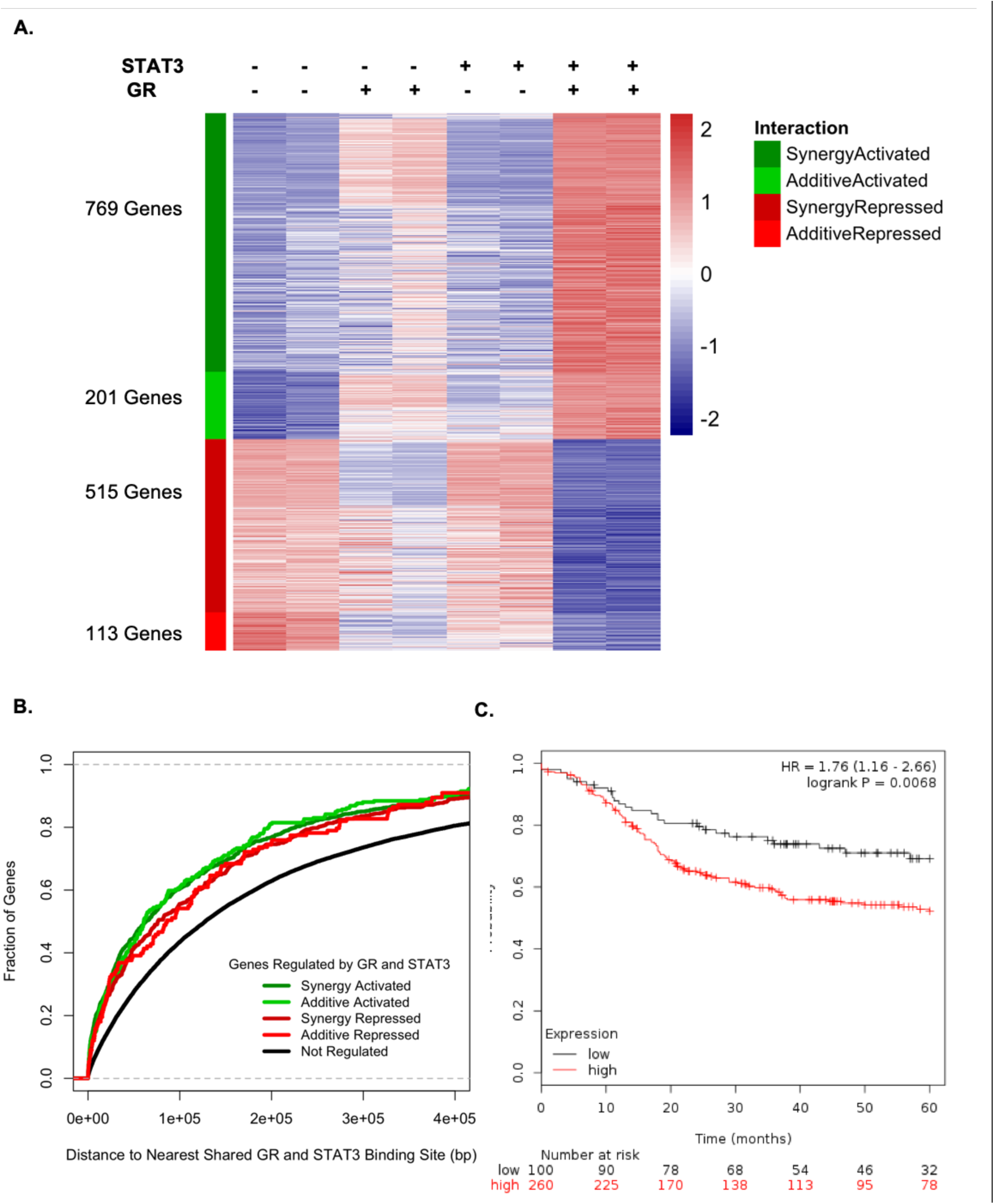
GR and STAT3 cooperate to regulate the expression of 1,598 genes. MDA-MB-231 cells were treated with dexamethasone to induce GR and SH4-54 to inhibit STAT3. **A.** A heatmap of genes that are regulated by both GR and STAT3 in a manner that fits additive or synergistic models for activation or repression (BH adjusted p < 0.05). **B.** Genes regulated by both GR and STAT3 are closer to shared GR and STAT3 binding sites than genes that are not regulated by these factors. **C.** Higher mean expression of genes that are upregulated by GR and STAT3 is associated with shorter relapse-free survival in basal-like breast cancer patients.

To determine if the genes that are cooperatively regulated by GR and STAT3 are near shared GR and STAT3 basal-like specific binding sites, the distance between the transcription start site of each gene and the nearest shared GR and STAT3 binding site was calculated. Consistent with previous observations, a larger fraction of genes that are cooperatively activated by GR and STAT3 are close to a shared site, when compared to repressed genes, or genes that did not change expression when these TFs were modulated (**Figure 6B**).

Together these results indicate that GR and STAT3 cooperate to synergistically regulate the expression of hundreds of genes.

### GR and STAT3 regulate the basal-like gene expression program

To test the hypothesis that GR and STAT3 are key regulators of the basal-like gene expression program, RNA-seq data from 28 breast cancer cell lines (18 basal-like and 10 luminal) were analyzed to identify genes that exhibit subtype-specific expression. Of the 2,485 genes that had significantly higher expression in basal-like breast cancer, 439 were significantly upregulated in HCC70 and MDA-MB-231 when comparing dexamethasone induction to baseline media, and standard media to STAT3 siRNA or SH4-54 inhibition of STAT3. These 439 basal-like genes that are up-regulated by GR and STAT3 have a significant overlap with genes that were previously characterized as more highly expressed in basal-like breast cancer compared to luminal (Hypergeometric FDR adjusted q-value = 2.6 × 10^-95^)(37). This result confirms that the GR and STAT3 regulated genes identified in this analysis have been associated with basal-like breast cancer in previous studies.

Further characterization of the 439 basal-like genes upregulated by GR and STAT3 revealed enrichment for cellular processes and pathways associated with the aggressive nature of basal-like breast cancer. These genes were significantly enriched for the Gene Ontology Biological Process “Positive Regulation of Cell Proliferation” (Hypergeometric FDR adjusted q-value 2.04 × 10^-17^) (38-40), “hallmarks that define the epithelial-mesenchymal transition” (Hypergeometric FDR adjusted q-value 7.22 × 10^-15^) (38-40), “mammary stem cells” (Hypergeometric FDR adjusted q-value = 1.04 × 10^-22^) (41), and “adult tissue stem cells” (Hypergeometric FDR adjusted q-value =2.13 × 10^-20^)(42).

Notably, of the 439 basal-like genes upregulated by GR and STAT3, 34 are TFs themselves, including those that are known to be upregulated by major signaling pathways involved in basal-like breast cancer growth including EGFR signaling (ELK3, ETS2, CEBPD, NFIL3, MBNL2, HIVEP2) (43), and TGFB signaling (ELK3, ETS2, SNAI1, MAFB)(44). Additionally, the genes regulated by GR and/or STAT3 include 5 TFs (ETV6, NFIL3, HIF1A, DR1, and TFCP2L1) that were previously identified as “embryonic stem cell” TFs that are preferentially and coordinately overexpressed in the high-grade, ER-negative breast cancer tumors(45). These results suggest GR and STAT3 are upstream of the key TFs that control growth and stemness features in basal-like TNBC. These 439 basal-like genes are upregulated by GR and STAT3 in both HCC70 and MDA-MB-231 cell lines. To determine if they are associated with prognosis in patients with basal-like breast cancer, Kaplan Meier survival plots were generated using public gene expression microarray data from 360 basal-like patient tumors for whom five-year relapse-free survival data were available. Patients who had higher mean expression of these genes in their tumors had significantly shorter relapse-free survival than patients with lower expression of these genes (Hazard Ratio =1.76, logrank p=0.0068) (**Figure 6C**). This result indicates that even amongst basal-like patients, higher expression of genes that are upregulated by GR and STAT3 is associated with a worse prognosis.

### Inhibiting both GR and STAT3 reduces cell proliferation synergistically

The observation that GR and STAT3 cooperate to regulate a large number of genes in the basal-like gene expression signature that are associated with cell proliferation and poor prognosis led us to investigate whether drugs that modulate GR and STAT3 would affect cell proliferation. To first evaluate the independent roles of GR and STAT3 in cell proliferation, the MDA-MB-231 cell line was treated with a range of doses of dexamethasone or the STAT3 inhibitor SH4-54 separately and was monitored for 72 hours on the Incucyte Zoom live cell imaging instrument. Reducing STAT3 by increasing doses of the STAT3 inhibitor SH4-54 resulted in decreased growth in a dose-dependent manner (**Figure 7A**). Reducing GR activity by decreasing amounts of dexamethasone resulted in decreased cell growth in a dose-dependent manner (**Figure 7B**). These results are consistent with previous publications that report that inhibiting GR or STAT3 separately decreases basal-like breast cancer proliferation (18,46-48).

**Figure 7.**
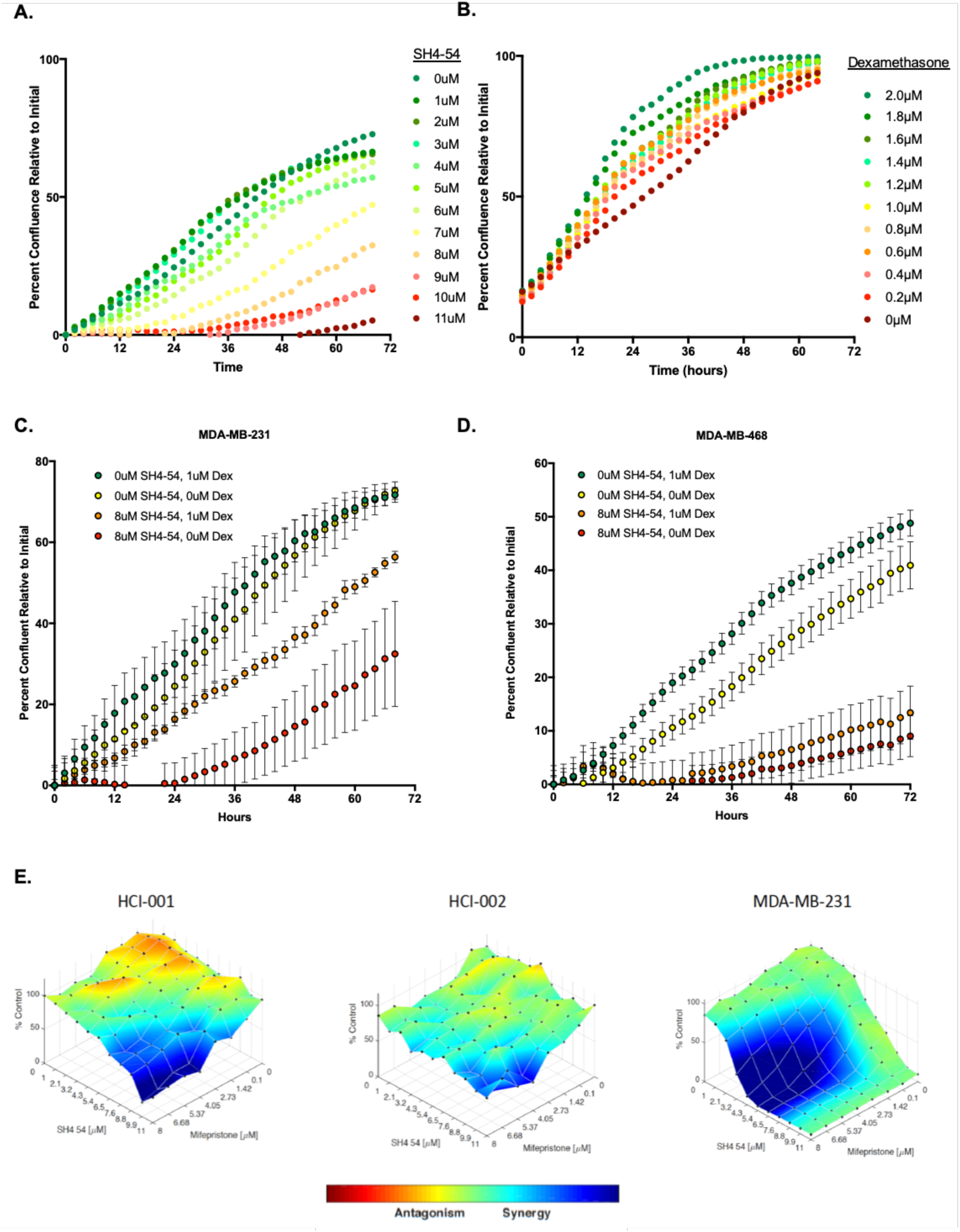
Inhibiting GR and STAT3 leads to decreased growth and proliferation in basal-like cell lines and patient-derived organoids. **A.** MDA-MB-231 cells treated with increasing doses of the STAT3 inhibitor SH4-54 have decreased growth rate in a dose-dependent manner. **B.** MDA-MB-231 cells treated with decreasing doses of dexamethasone have decreased growth rate in a dose-dependent manner. Inhibiting both GR and STAT3 in MDA-MB-231 (**C**) and MDA-MB-468 (**D**) cell lines leads to decreased growth compared to inhibiting either factor alone. **E.** Synergistic decreases in cell proliferation are observed after 72 hours when patient-derived organoid cultures (HCI-001 and HCI-002) and MDA-MB-231 cell lines are treated with increasing doses of both the STAT3 inhibitor SH4-54 and the GR inhibitor mifepristone.

To evaluate whether modulating both TFs simultaneously leads to differences in proliferation, the MDA-MB-231 and MDA-MB-468 cell lines were treated with different combinations of dexamethasone and SH4-54. The combination of inhibiting GR by withholding dexamethasone, and inhibiting STAT3 with 8µM SH4-54 resulted in the slowest cell proliferation (**Figure 7C & 7D**). Conversely, the activation of GR with 1µM dexamethasone combined with the endogenous activity of STAT3 resulted in the fastest cell proliferation (**Figure 7C & 7D**). Notably, when either GR or STAT3 were active alone, this led to intermediate proliferation rates. This result supports the hypothesis that GR and STAT3 cooperate to increase cell proliferation, and dual inhibition of both GR and STAT3 results in slower cell proliferation than inhibiting either factor alone.

To expand this analysis beyond cell lines, proliferation of patient-derived organoid cultures was analyzed in the context of GR and STAT3 inhibition. Two patient-derived organoid cultures previously derived from primary basal-like TNBC tumors (HCI-001 and HCI-002) were utilized for this experiment(49,50). The standard media for the patient-derived organoid cultures includes 1 µg/ml hydrocortisone, which is a glucocorticoid that induces GR activity. Therefore, escalating doses of mifepristone were used to inhibit GR activity. Mifepristone also inhibits progesterone receptor (PR) activity, but basal-like cells do not express PR, and thus this is not a confounding factor in these experiments. The STAT3 inhibitor SH4-54 was used in escalating doses to inhibit STAT3 activity. The cell line MDA-MB-231 was included in this experiment as a positive control and was grown in the same media as the organoids. The proliferation rate of the organoids is much slower than the cell line, which resulted in a narrower dynamic range for observing changes in proliferation. Despite this technical limitation, inhibition of GR and STAT3 for 72 hours resulted in significant decreases in cell proliferation in a dose-dependent manner (**Figure 7E**). An analysis of the relative decrease in cell proliferation across drug doses indicates that inhibiting both GR and STAT3 provides synergistic, rather than additive, decreases in cell proliferation in the MDA-MB-231 cell line, as well as both patient-derived organoid cultures (**Figure 7E**). This result confirms that inhibiting the activity of both GR and STAT3 reduces cell proliferation in basal-like breast cancer patient-derived organoids.

## Discussion

The goal of this study was to determine which TFs drive the basal-like TNBC gene expression program. Genome-wide analysis of DNA methylation, chromatin accessibility, TF binding, and gene expression indicate that GR and STAT3 binding is enriched at genomic regulatory regions that are specifically open in basal-like TNBC. These TFs bind to the same regulatory regions and cooperate to regulate hundreds of genes in the basal-like expression signature, including many other TFs. The genes that are cooperatively up-regulated by GR and STAT3 are transducers of major signaling pathways implicated in basal-like breast cancer growth (EGFR and TGFB), are associated with aggressive features of basal-like breast cancer (proliferation, stemness and epithelial to mesenchymal transition), and are associated with shorter relapse-free survival in patients. Furthermore, inhibiting both GR and STAT3 leads to synergistic reductions in cell proliferation in multiple cell lines and patient-derived organoid cultures. Together these data suggest that inhibiting both GR and STAT3 simultaneously can reduce expression of the basal-like gene expression signature and create a synthetic lethality in TNBC that could be exploited therapeutically.

While GR and STAT3 have been separately implicated in regulating gene expression in basal-like breast cancer previously(18,48,51,52), their cooperative role has not been previously described. This study makes it apparent that understanding this cooperation is crucial to investigating their roles in driving basal-like TNBC gene expression and phenotypes, as well as arriving at more effective therapeutic strategies. Inhibiting GR alone is not effective; basal-like cell lines proliferate and express some components of the basal-like gene signature without glucocorticoids in their media. Inhibiting STAT3 alone is not completely effective; previous studies have shown that SH4-54 killed basal-like cell lines in culture, but was insufficient to completely stop tumor growth in vivo where glucocorticoids are naturally present (24). These results indicate that inhibiting either of these TFs alone is not sufficient to abrogate the pro-proliferative gene expression program of basal-like breast cancer in vivo, but this study suggests that inhibiting both GR and STAT3 could be a novel and effective therapeutic strategy.

The study of GR’s role in basal-like breast cancer has important clinical implications because potent synthetic glucocorticoids are often prescribed to breast cancer patients undergoing chemotherapy to prevent side-effects such as nausea, loss of appetite, and rare severe immune reactions. Our preliminary data supports a handful of other studies that report that induction of GR with glucocorticoids could promote tumor growth and therapy resistance (15,17,18,52). In our study specifically, it is clear that glucocorticoid induction of GR leads to increased cell proliferation in basal-like breast cancer cell lines and patient-derived organoids. It also leads to increased expression of genes that are cooperatively regulated by STAT3, a known promoter of cancer growth and invasion (53-58). Recently, clinical trials have been initiated to test whether inhibiting GR directly with mifepristone or by inhibiting its chaperone, Hsp90, will enhance the efficacy of chemotherapy in basal-like breast cancer (16,59). Our results support this line of investigation and suggest that combination therapy that inhibits both GR and STAT3 could be even more effective in treating this disease.

## Acknowledgements

This work was supported by American Cancer Society Research Scholar Grant Award # 132596-RSG-18-197-01-DMC awarded to KEV. We acknowledge support of funds in conjunction with grant P30 CA042014 awarded to Huntsman Cancer Institute and to the Nuclear Control Program at Huntsman Cancer Institute. Research reported in this publication utilized the High-Throughput Genomics and Bioinformatic Analysis Shared Resource at Huntsman Cancer Institute at the University of Utah and was supported by the National Cancer Institute of the National Institutes of Health under Award Number P30 CA042014. This research was also supported by funding from the HudsonAlpha Institute, a Cancer Research Fund from the State of Alabama to HudsonAlpha Institute, Susan G. Komen, and the NIH-National Cancer Institute Comprehensive Cancer Center Core Support Grant (5P30CA013148). JMM was supported by a fellowship from the National Science Foundation Graduate Research Fellowship Program (NSF-GRFP).

## Author Contributions

MEC, JMM, JMG, KPG, PGO, and SLP performed experiments. MEC and KEV performed analysis and wrote the manuscript. PY and JT provided reagents. DJB, BEW, and RMM provided expertise and feedback.

## Materials and Methods

### Cell Lines

The following basal-like breast cancer cell lines were used for this study: HCC1569, HCC1954, BT-20, HCC1143, HCC1187, HCC1599, HCC1937, HCC70, MDA-MB-468, 2-LMP, BT-549, HCC38, MDA-MB-157, MDA-MB-231, MDA-MB-436, SUM102, SUM149, SUM159.

The following luminal breast cancer cell lines were used for this study: DY36T2, ZR-75-30, MDA-MB-453, SK-BR-3, BT-474, MDA-MB-361, MCF-7, MDA-MB-134, T-47D, ZR-75-1.

Cells were obtained and cultured as described previously(60). Cell lines passaged for more than 3 months were authenticated every 6 months using Short Tandem Repeat Testing service offered through the University of Utah DNA Sequencing Core Facility.

### Reduced Representation Bisulfite Sequencing (RRBS)

RRBS and primary analysis of CG methylation was performed on the panel of cell lines using the method described previously(61). CG positions with at least 10x coverage in at least 80% of the cell lines were used for subsequent analysis. To identify CGs that were differentially methylated between basal-like and luminal cell lines, Linear Regression was performed in the R software package version 3.5.0 using the lm function. P-values were adjusted for multiple hypothesis testing using the p.adjust function in R and the Benjamini-Hochberg method. CGs with adjusted p-values less than 0.05 were visualized in R using the pheatmap package version 1.0.10.

### Enrichment of ENCODE Transcription Factor Binding Sites

A file named wgEncodeRegTfbsClusteredV2.bed.gz containing TF binding sites from ChIP-seq experiments performed on 149 TFs by the ENCODE Project(26) in a variety of cell lines were downloaded from the UCSC genome browser(62). Bedtools intersect (63) was used to determine which CG positions in the RRBS dataset overlapped binding site positions for each TF in the file. The number of overlaps with basal-like and luminal specific CGs was normalized to the number of overlaps to all CGs in RRBS to calculate a fold-enrichment.

### Analysis of TCGA RNA-seq and ATAC-seq

RNA-seq counts file for TCGA Breast Cancer samples were downloaded from the UCSC Xena browser(64,65). These files contained log counts, so they were transformed to counts in R version 3.5.0. DESEQ2 version 1.20.0 (66) was used to identify genes that were differentially expressed between basal-like and luminal tumors using the PAM50 Subtype classification provided by TCGA and available for download through the UCSC Xena browser(64,65). Basal specific genes were identified using an adjusted p-value < 0.05 and coefficients less than 0. Luminal specific genes were identified using a p-value < 0.05 and coefficients greater than 0.

ATAC-seq data from TCGA breast tumors were downloaded as counts files and bigwig files from the supplementary material provided in the recent publication by Corces et al. (29). DESEQ2 version 1.20.0 (66) was used to identify peaks that were differentially accessible between basal-like and luminal tumors using the PAM50 Subtype classification provided by TCGA and available for download through the UCSC Xena browser(64,65). Pheatmap package version 1.0.10 was used to visualize basal-specific peaks that were identified using an adjusted p-value < 0.05 and coefficients less than 0. The ATAC-seq peaks specifically open in basal-like tumors were analyzed for enrichment of motifs for JUN (MA0491.1), STAT3 (MA0144.1), and GR(MA0113.2) downloaded from JASPAR2018(67). AME(68) was used to perform the analysis of sequences in ATAC-seq peaks compared to a primary sequence shuffled conserving 2-mer frequencies, and p-values were calculated using Fisher’s exact test.

To analyze ATAC-seq signal from TCGA breast tumors over the basal-specific GR and STAT3 shared binding sites, the UCSC hgLiftOver was used to convert the GR and STAT3 shared sites bed file to the hg38 genome build to be compatible with TCGA ATAC-seq alignments. Deeptools version 3.1.0(69) computeMatrix was used to generate a matrix of counts over the GR and STAT3 shared sites for each sample. The mean accessibility was computed across the regions for each sample using R version 3.5.0. The Mann Whitney test was used to determine if the average normalized ATAC-seq read depth across regions was significantly different between basal-like and luminal tumors. The mean and standard deviation of the average normalized depth across basal-like and luminal samples was graphed using GraphPad Prism 7.0c.

Violin plots for averaged bigwig read counts from basal-like and luminal tumors at GR and STAT3 shared sites were created with vioplot_0.3.2, using default parameters and areaEqual=T, h=0.04, wex=0.9.

### ChIP-seq

ChIP-seq for GR and STAT3 was performed on 4 basal-like cell lines (SUM159, MDA-MB-231, HCC1937, and HCC70) and 4 luminal cell lines (MDA-MB-361, BT-474, MDA-MB-453, and MCF-7).

For ChIP experiments, protein-DNA complexes were covalently cross-linked by incubating cells in 1% formaldehyde for 10 min at room temperature. Cells were incubated with 0.125 M glycine for 5 min, to quench cross-linking reaction. Cells were washed and scraped with PBS (pH 7.4) (Lonza). Cells were lysed with Farnham Lysis Buffer (5mM PIPES at pH 8.0, 85 mM KCl, 0.5% NP-40) containing protease inhibitor (Roche). Cell lysate was centrifuged at 2,000 rpm for 5 min at 4 °C. The crude nuclear extract contained in the supernatant was stored at −80 °C. ChIP-seq was performed as previously described(70) using antibodies for GR (sc-1003, Santa Cruz Biotechnology) and STAT3 (sc-482, Santa Cruz Biotechnology). A thorough version of the ChIP-seq protocol used in this study is available on the ENCODE Project website:

https://www.encodeproject.org/documents/df9dd0ec-c1cf-4391-a745-a933ab1af7a7/@@download/attachment/Myers_Lab_ChIP-seq_Protocol_v042211.pdf

STAT3 ChIP-seq datasets have been deposited in the NCBI Gene Expression Omnibus (GEO) accession number GSE85579.

Fastq files from ChIP-seq were aligned to the hg19 build of the human genome using Bowtie with the following parameters: -m 1 -t –best -q -S -l 32 -e 80 -n 2. ChIP-seq peaks were identified by comparing GR ChIP-seq in cells induced with dexamethasone to GR ChIP-seq in cells treated with ethanol, and STAT3 ChIP-seq to input control libraries. Peaks were called using Model-Based Analysis of ChIP-seq-2 (MACS2) (71) with a p value cutoff of 1e -10 and the mfold parameter constrained between 15 and 100. Bedtools merge (63) was used to merge bed files of MACS2 narrow peak calls from each of the ChIP-seq experiments. Bedtools coverageBed (63) was used to extract read counts under each merged peak in each ChIP-seq experiment in each cell line. DESEQ2 version 1.20.0 (66) was used to identify peaks with significantly different read depth between basal-like and luminal cell lines (adjusted p-value < 0.05), and significantly different read depth between GR and STAT3 ChIP-seq experiments (adjusted p-value < 0.05). A multivariate model was used to identify shared GR and STAT3 peaks whose read depth was significantly different between subtypes in both GR and STAT3 ChIP-seq experiments (adjusted p-value < 0.05), but not significantly different between GR and STAT3 in the same subtype (adjusted p-value > 0.05). Pheatmap package version 1.0.10 and Deeptools version 3.1.0(69) computeMatrix and plotHeatmap functions were used to create the heatmaps of ChIP-seq data.

### RNA-seq

RNA-seq of the panel of cell lines was described previously (72) and the data are publicly available through NCBI Gene Expression Omnibus (GEO) accession number GSE58135.

RNA-seq was also performed on MDA-MB-231 and HCC70 cell lines treated alone or in combination with dexamethasone, STAT3 siRNA, and SH4-54 as follows:

STAT3 siRNA knockdown utilized the ON-TARGETplus Human STAT3 siRNA kit from GEHealthcare (L-003544-00-0005, GE Healthcare). This SMARTpool siRNA contains four pooled siRNAs, each targeting a separate region of the STAT3 RNA sequence. ON-TARGETplus Non-targeting siRNA #1 (D-001810-01-05) as a non-targeting control. The siRNA SMARTpool and Non-targeting siRNA target sequences are below:

ON-TARGETplus SMARTpool siRNA J-003544-07, STAT3: GAGAUUGACCAGCAGUAUA

ON-TARGETplus SMARTpool siRNA J-003544-08, STAT3: CAACAUGUCAUUUGCUGAA

ON-TARGETplus SMARTpool siRNA J-003544-09, STAT3: CCAACAAUCCCAAGAAUGU

ON-TARGETplus SMARTpool siRNA J-003544-10, STAT3: CAACAGAUUGCCUGCAUUG

ON-TARGETplus Non-targeting siRNA #1: UGGUUUACAUGUCGACUAA

The siRNA transfection experiments were performed in 6-well plates in triplicate. Lipofectamine RNAiMAX Transfection Reagent (ThermoFisher Scientific) was used per manufacturer’s instructions. To each well containing cells, 250 μl siRNA-transfection reagent mix was added to each well, for a final concentration of 25 pmol siRNA in 7.5 μl Lipofectamine RNAiMAX reagent per well.

The dexamethasone treatments were performed in 6-well plates in triplicate. Cell were treated with 100 nM dexamethasone (D4902, Sigma Aldrich) or an equal volume of 100% molecular biology grade ethanol vehicle control (E7023, Sigma Aldrich) for 4 hours. Cells were lysed with Buffer RL (Norgen Biotek) containing 10% beta-mercaptoethanol. Total RNA was extracted using the Animal Tissue RNA Purification Kit (Norgen Biotek). RNA-seq libraries for the STAT3 siRNA and Dexamethasone treatment were prepared from 250 ng total RNA via polyA-selection (Dynabead mRNA Purification Kit, Invitrogen) followed by transposase-mediated non-stranded library construction(73). Libraries were pooled and sequenced on an Illumina HiSeq 2000 or HiSeq 2500 sequencer using paired-end 50bp reads with a 6 bp index read. TopHat v1.4.1 was used to align RNA-seq paired reads to GENCODE version 9. Cufflinks v1.3.0 and BEDTools were used to calculate raw counts for each GENCODE transcript.

DESEQ2 version 1.20.0 (66) was used to identify genes with significantly different expression between treatment and control conditions in each cell line separately. To visualize differential expression of genes across cell lines with different base-line expression, gene expression was divided by the mean in the control sample, and normalized values were displayed in a heatmap using Pheatmap package version 1.0.10 in R.

For the GR and STAT3 interaction studies, MDA-MB-231 cells were plated in 6-well plates at ∼300,000 cells in 4ml of media per well and grown for 24 hours. Three replicates wells were treated for 24 hours with 1µM of Dexamethasone (DEX), 8µM SH4-54, no treatment, or combination of DEX and SH4-54. The cells were lysed using 350µL of Qiagen RLT buffer and 1% BME. RNA was extracted using Norgen Animal Tissue RNA Purification Kit according to manufacturer instructions. RNA-seq libraries were prepared using Illumina TruSeq Stranded RNA Kit with Ribo-Zero Gold and sequenced on the Illumina HiSeq 2500 as 50 cycle single-end reads. Reads were mapped using HISAT2(74) version 2.1.0 with parameters --dta -p 28 -t -q -U and sorted with samtools(75). A counts file was generated from the bam files using featureCounts version 1.5.1(76).

To identify genes that were additively and synergistically regulated by both GR and STAT3 the linear model (lm) function in R was used. DESEQ2 normalized counts were fit to a multivariate model that included a term for GR activity, a term for STAT3 activity, and an interaction term (GR:STAT3). Coefficients and p-values for each gene were used to identify those with additive and synergistic positive and negative regulation by GR and STAT3. Synergy was defined as a significant p-value (<0.05) for the interaction term, and non-significant p-values (>0.05) for the individual GR and STAT3 terms. Additivity was defined as a non-significant p-value for the interaction term, and significant p-values for both of the individual GR and STAT3 terms with coefficients in the same direction. Coefficients from the linear model were used to determine whether the interactions between GR and STAT3 led to increased or decreased gene expression.

RNA-seq data from STAT3 siRNA treatments are publicly available through NCBI Gene Expression Omnibus (GEO) accession number: GSE85579.

### Kaplan Meier analysis

The Kaplan-Meier Plotter tool (http://kmplot.com)(32) was used to analyze the association between gene expression and relapse-free survival. A Kaplan Meier survival plot was generated using public gene expression microarray data from patient tumors for whom relapse-free survival data were available. The intrinsic subtype classification provided by the Kaplan-Meier Plotter tool was used to select basal-like and luminal cases for analysis (77). The analysis was performed for GR, STAT3 and 439 basal-like genes that are upregulated by GR and STAT3. The following selections were applied to all analyses: only one JetSet best probe (78) for each gene was used, multigene signatures used the mean expression of the selected probes, relapse-free survival was selected for the analysis, patients were censored at the follow-up threshold (60 or 120 months), biased arrays were excluded, and redundant samples were removed. The most significant cutpoint was used to split patients into two groups (“autoselect best cutoff” option).

### Live cell imaging of proliferation

Proliferation assays were performed using the IncuCyte ZOOM live cell imaging system (Essen BioSciences). MDA-MB-231 and MDA-MB-436 cells were plated at 20,000 cells per well in 96 well plates. For titration experiments, cells were treated a range of doses of SH4-54 between 0uM and 11uM of SH4-54, or a range of doses of dexamethasone between 0uM and 2uM. For combination treatments cells were treated with 8uM SH4-54 and 1uM dexamethasone, alone and in combination. Plates were imaged every 2 hours for 72 hours, and cell confluence was calculated by the IncuCyte ZOOM software. Relative percent confluence was calculated by subtracting the percent confluence at each timepoint by the starting percent confluence at the first timepoint.

### Patient derived organoid proliferation assay

Two patient derived organoid cultures previously derived from primary basal-like TNBC tumors (HCI-001 and HCI-002) were utilized for this experiment(49,50). The organoids were cultured in media composed of Advanced DMEM/F12, (Thermo Fisher 12634028), 5% FBS ThermoFisher 26140079), 1x Hepes (ThermoFisher 15630080), 1x GlutaMAX Supplement (Thermo Fisher 35050061), 1 ug/ml Hydrocortisone (Sigma Aldrich H0888), 10 ng/ml hEGF (Sigma Aldrich E9644), 50 ug/ml Gentamicin (Genesee 25-533), 10 uM Y-27632 (Selleckchem S1049). Organoids were seeded as 50-100 organoids in 5% Matrigel (Corning 354230) per well of a 384 well plate, as described previously(49). MDA-MB-231 cell lines were included as a positive control in this experiment and plated at 500 cells per well in a 384 well plate. Cells were treated with all possible combinations of titrated doses of SH4-54 (0, 1, 2.1, 3.2, 4.3, 5.4, 6.5, 7.6, 8.8, 9.9, and 11uM) and mifepristone (0, 0.1, 1.42, 2.73, 4.05, 5.37, 6.68, and 8uM) in combination. Cell viability was measured relative to untreated cells after 72 hours using the CellTiter Glo 3D (Promega) that measures ATP. Lowe additivity analysis was used to compute synergy scores(79).

